## Extended data set for "Pseudoirreversible inhibition elicits persistent efficacy of a sphigosine-1-phosphate receptor-1 antagonist"

**TITLE:**

Contact Information: Taishin Akiyama, Ph.D.

Postal address: 1-7-22 Suehiro-cho, Tsurumi-ku, Yokohama 230-0045, Japan

Extended Data Figure 1

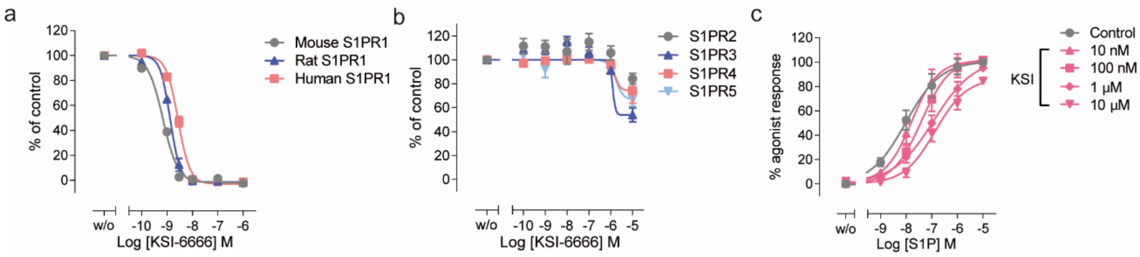


**Extended Data Figure 1.** $Ca^{2+}$ mobilization assay for KSI-6666.

**a** and **b**. Inhibitory effect of KSI-6666 on $Ca^{2+}$ mobilization induced by S1P receptor family signaling. $Ca^{2+}$ mobilization was induced by 10 nM Merck S1PR1 agonist for S1PR1 of human, mouse, or rat (a), or by 100 nM S1P for human S1PR2, S1PR3, S1PR4, or S1PR5 (b) in CHO-K1 cells expressing S1P receptors. Gα15 was co-expressed with S1PR1, S1PR4, or S1PR5. Data are the mean ± s.e.m. of three independent experiments.

**c**. Shift in $EC_{50}$ for S1P activation of human S1PR1 by KSI-6666 showing its competitive antagonistic activity. CHO-K1 cells expressing human S1PR1 and Gα15 were co-treated with mixtures of serial dilutions of S1P and KSI-6666. The increase in cellular calcium level induced by 10 μM S1P alone was calculated as 100% response and that in conditions without agonist or the test compound was defined as 0%. Nonlinear regression analysis of the concentration–response curve was performed in GraphPad Prism 6 software. Data are the mean ± s.e.m. of four independent experiments.

Extended Data Figure 2

a

| Test article | Control | | | | | KSI-6666 | | | | | KSI-6666 | | | | | KSI-6666 | | | | | FTY720 | | | | |
|---|---|---|---|---|---|---|---|---|---|---|---|---|---|---|---|---|---|---|---|---|---|---|---|---|---|
| Dose | NA | | | | | 0.05 mg/kg b.i.d. | | | | | 0.5 mg/kg b.i.d. | | | | | 5 mg/kg b.i.d. | | | | | 5 mg/kg q.d. | | | | |
| Animal No. | 1 | 2 | 3 | 4 | 5 | 6 | 7 | 8 | 9 | 10 | 11 | 12 | 13 | 14 | 15 | 16 | 17 | 18 | 19 | 20 | 21 | 22 | 23 | 24 | 25 |
| Findings | N | N | N | N | N | N | N | N | N | Y | N | N | Y | Y | Y | Y | Y | N | Y | Y | N | Y | N | N | N |
| Infiltration, macrophage | | | | | | | | | | - | | | ± | 1+ | 1+ | 1+ | ± | | 1+ | 1+ | | ± | | | |
| Edema, perivascular | | | | | | | | | | ± | | | - | - | ± | - | - | | - | - | | - | | | |

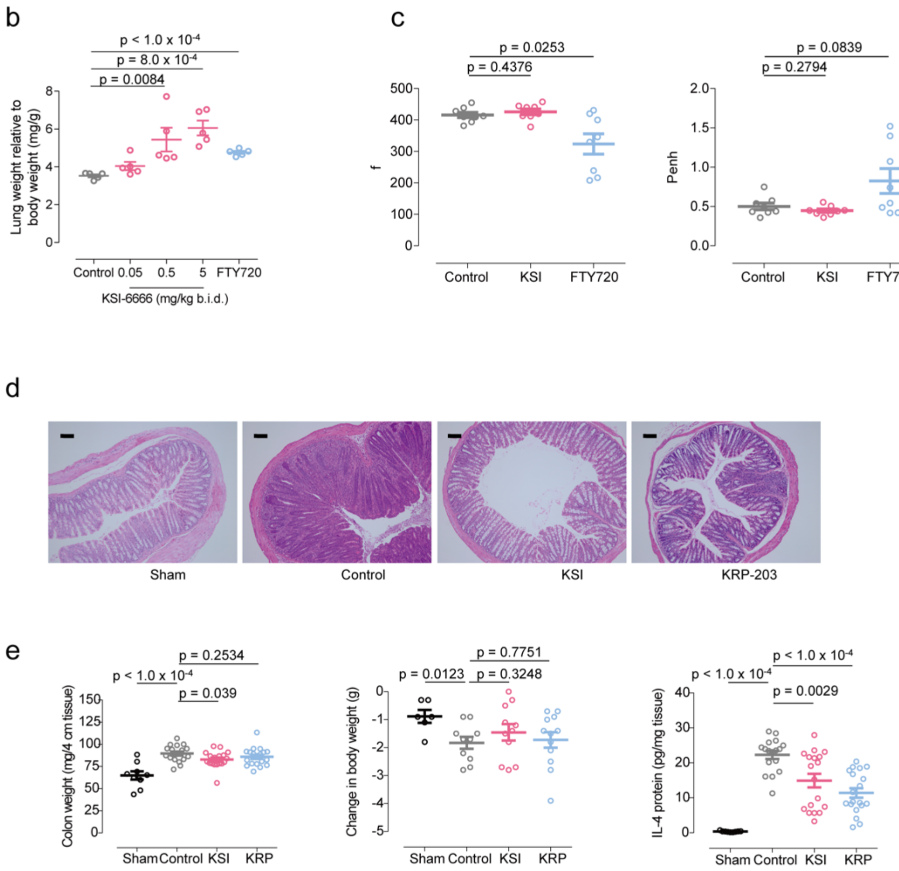


**Extended Data Figure 2. In vivo effect of KSI-6666 administration on normal mouse and mouse disease model.**

**a.** Histological findings in lungs of rats after repetitive oral administration of test compounds for 4 weeks. The test compound dissolved or suspended in 0.5% methyl cellulose 400 solution was orally administered to 8-week-old male Wistar Hannover rats for 4 weeks. KSI-6666 was administered at 0.05, 0.5, and 5 mg/kg b.i.d., and FTY720 at 5 mg/kg q.d. Following euthanasia, lungs were collected and histologically examined ($n$ = 5 /group). N: no finding; Y: finding present; −: not remarkable/not present; ±: minimal; 1+: slight; 2+: Moderate; 3+: Marked.

**b.** Lung weight of rats after repetitive oral administration of test compounds for 4 weeks. The test compound dissolved or suspended in 0.5% methyl cellulose 400 solution (Wako) was orally administered to 8-week-old male Wistar Hannover rats (Nihon Clinical Science Laboratory) for 4 weeks. KSI-6666 was administered at 0.05, 0.5, and 5 mg/kg b.i.d., and FTY720 at 5 mg/kg q.d. Following euthanasia, lungs were collected and weighed. Statistical analysis was performed by Student's t-test for FTY720, and by one-way ANOVA followed by Dunnett's test for KSI-6666 (mean ± s.e.m., $n = 5$ /group).

**c.** Compound-induced change to respiratory function in mice. The test compound dissolved or suspended in 0.5% methyl cellulose 400 solution was orally administered to 8-week-old male BALB/cCr mice (Japan SLC). Two equimolar amount of hydrochloric acid (Wako) was mixed only in the suspension of KSI-6666. KSI-6666 was administered at 20 mg/kg and FTY720 at 5 mg/kg. Four hours after administration, spontaneously breathing awake mice unrestrained were placed in the whole-body plethysmographic chamber (Buxco) connected to a pressure transducer. Acclimatization of mice to the chamber was performed on the day before the assessment for 20 min and on the day of assessment for 17 min. Respiratory rate (f: breaths per minutes) and respiratory pressure curves (Penh) were recorded for 3 min by FinePointe Systems (Buxco). Statistical analysis was performed by either Student's t-test or Welch's t-test according to F-test results (mean ± s.e.m., $n = 8$ /group).

**d.** Representative histological images of the T cell transfer colitis model were exhibited (scale bar = 100 μm).

**e.** Efficacies of KSI-6666 (30 mg/kg/total) and KRP-203 (3 mg/kg/total) in oxazolone-induced colitis model. A total of 150 μL of 3% 4-ethoxymethylene-2-phenyl-2-oxazolin-5-one (oxazolone; Sigma-Aldrich) dissolved in 100% ethanol (Wako) was applied on shaved back skins of 9–10-week-old male BALB/cCr mice for sensitization. One week after sensitization, 100 μL of 1% oxazolone dissolved in 50% ethanol oxazolone was intrarectally administered to the mice. On the same day, test compounds suspended in the vehicle consisting of a 1:9 mixture of 0.5% methyl cellulose 400 solution (Wako) and distilled water (Otsuka) were orally administered to mice twice (before and after sensitization). Mice were sacrificed 24 h after sensitization, and colons were collected and weighed. Homogenates of colons were prepared using TisseLyser (QIAGEN) and the amount of interleukin (IL)-4 protein was evaluated using a Mouse IL-4 Quantikine ELISA Kit (R&D Systems). Change in body weight was calculated by the difference between the day of intrarectal oxazolone administration and the end of study.

Statistical analysis was performed by either Student's t-test or Welch's t-test according to F-test results (mean ± s.e.m., $n$ = 9–19).

### Extended Data Figure 3

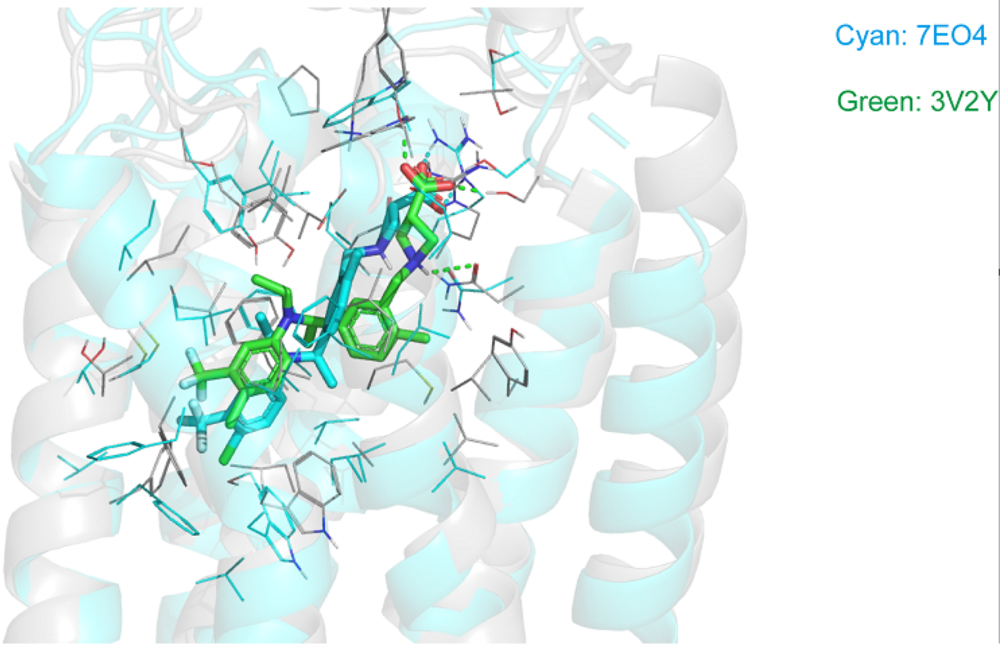


**Extended Data Figure 3. Comparison of *(R)*-KSI-6666 docking poses modeled from two different S1PR1 structures.**

Green indicates the structure obtained by induced fit docking protocol using an X-ray structure of W146-bound S1PR1(3V2Y). Cyan indicates that using an X-ray structure of siponimod-bound S1PR1 (7EO4).

Extended Data Figure 4

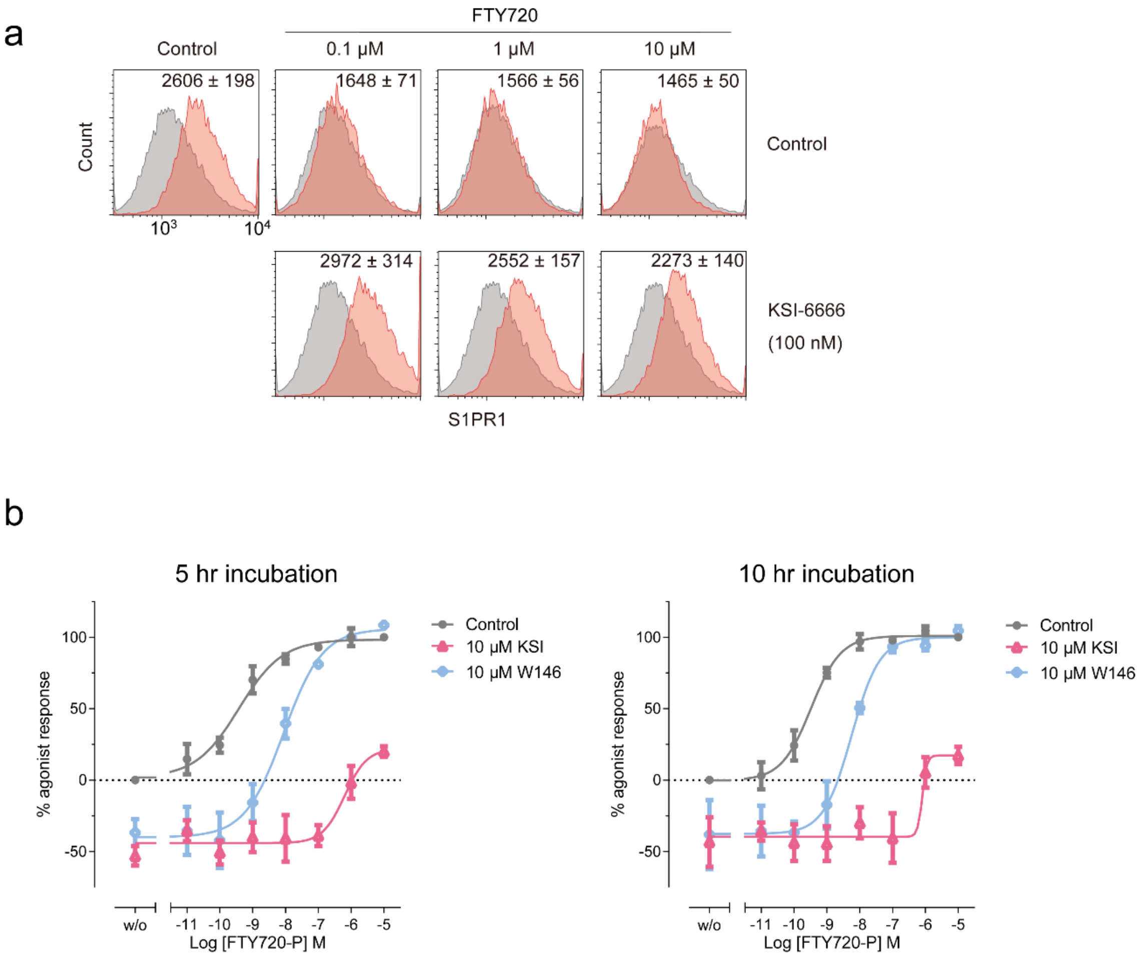


**Extended Data Figure 4. Inhibition of ligan-inducing internalization of S1PR1 by the addition of KSI-6666**

**a.** The inhibitory effect of KSI-6666 on FTY720-P induced receptor internalization evaluated by flow cytometry. KSI-6666 (100 nM) was preincubated with HEK293 stably expressing human S1PR1 in culture medium consisting of Dulbecco's Modified Eagle's Medium (Wako) supplemented with 10% fetal bovine serum (Biological Industries) for 15 min at 37°C, followed by the induction of receptor internalization by the addition of serial dilutions of FTY720-P (Cayman). Following 60-min incubation at 37°C, cells were washed by phosphate-buffered saline (Wako) and detached using Versene (Thermo Fisher Scientific). Cells were stained with anti-human S1PR1-PE antibody (R&D Systems FAB2016P) or isotype control antibody (BioLegend 401208). Dead cells were excluded via 7-aminoactinomycin D (BioLegend 420403) staining. Flow cytometric analysis was performed by spectral cell analyzer SA3800

(SONY), and data were analyzed by Flowjo software (BD). Data was obtained from three independent experiments and representative results are shown with the corresponding mean fluorescence intensity of all experiments (mean ± s.e.m.).

**b.** Functional reversibility of KSI-6666 and W146 examined after long incubation with compounds. HEK293 cells expressing HiBiT-tagged human S1PR1 were treated with KSI-6666 (10 μM) or W146 (10 μM) for 5–10 hours and concentration-dependent S1PR1 internalization induced by FTY720-P, represented by luciferase activity, was measured. Data was obtained from three independent experiments and results are expressed as the mean ± s.e.m.

Extended Data Figure 5

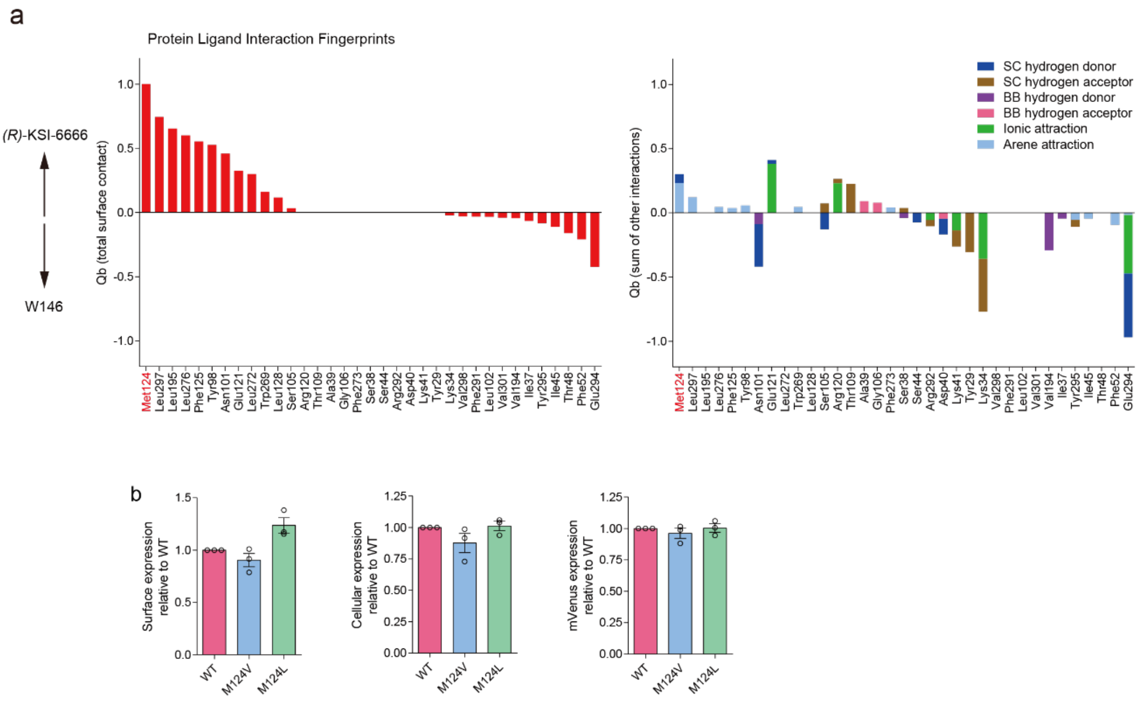


**Extended Data Figure 5. Protein-ligand interaction fingerprint analysis and expression levels of HiBiT-tagged wild-type and mutant human S1PR1.**

**a**. Significant protein–ligand interactions of *(R)*-KSI-6666 relative to W146 in protein–ligand interaction fingerprint (PLIF) analysis. The interaction between the compound and S1PR1 during MetaD simulation was represented in a fingerprint scheme by the PLIF tool. The interaction between the compound and each amino acid residue of S1PR1 was evaluated in two main categories: potential-energy based contacts and surface contacts. When an interaction above a threshold was observed for the residue, a fingerprint bit representing the interaction was assigned. Hydrogen bonding, ionic interaction, and π interaction (arene attraction) were identified as potential contacts with energies >0.5 kcal/mol and assigned as fingerprint bits in this analysis. For hydrogen bonding, each amino acid residue was broken down into side chain (SC) and backbone (BB) categories, and each was classified as either electron donor or acceptor. When a surface contact with area >15 Å$^2$ was observed, a fingerprint bit was assigned. The probability of each bit occurring for each residue for *(R)*-KSI-6666 compared with W146 was calculated as Qb, where Qb takes values from −1 to 1. A higher positive value indicates a higher probability of the bit occurring for *(R)*-KSI-6666. The columns show the value of Qb for each residue. The fingerprint bits shown are total surface contacts (left); and arene attraction (representing π interaction), ionic attraction, and BB hydrogen acceptor, BB hydrogen donor, SC hydrogen acceptor, and SC hydrogen donor interactions with *(R)*-KSI-6666 (right, different interactions shown in different colors).

**b-d**. Surface and cellular expression levels of HiBiT-tagged wild-type and mutant human S1PR1. HEK293 cells expressing HiBiT-tagged human S1PR1 (wild-type, or Val124 or Leu124 mutant) with mVenus were subjected to extracellular monitoring of HiBiT by using a Nano Glo HiBiT Extracellular Detection System (Promega) (**b**), or cellular monitoring of HiBiT by using a Nano Glo HiBiT Lytic Detection System (Promega) (**c**). The fluorescence intensity of mVenus, expressed as an internal control in each cell type, was evaluated (**d**). Data are the mean ± s.e.m. of three independent experiments.

**Supplementary Video 1**

A typical movie of MetaD simulation for the interaction between *(R)*-KSI-6666 and S1PR1

**Supplementary Video 2**

A typical movie of MetaD simulation for the interaction between W146 and S1PR1

**Supplementary Video 3**

A typical movie of MetaD simulation for the interaction between *(R)*-KSI-6666 and S1PR1 Val124 mutant

**SUPPLEMENTARY METHODS**

All reagents and solvents were purchased from commercial sources and used without further purification. All moisture and air sensitive reactions were carried out under an argon atmosphere. Reactions were monitored by thin layer chromatography (TLC) using Merck Kieselgel 60 F254 plates. Flash column chromatography was performed on YAMAZEN amino packed inject column and Biotage prepacked columns using an automated flash chromatography system (Biotage, Isolera One). $^{1}$H NMR spectra was recorded on a Bruker AV400 M spectrometer at 400.1 MHz. Chemical shifts (δ) were reported in parts per million (ppm) units using tetramethylsilane as an internal standard. Data were presented as follows; chemical shift, integration, multiplicity (s, singlet; d, doublet; t, triplet; q, quartet; m, multiplet; br, broad), and coupling constant. $^{13}$C NMR spectra were recorded on a Bruker AV400M spectrometer at 100.6 MHz with complete proton decoupling. Chemical shifts (δ) were reported in parts per million (ppm) with the solvent as the internal reference (δ 77.16 in $CDCl_3$). High resolution mass spectra (HRMS) was recorded on an Agilent Technologies 6520 Accurate-Mass Q-TOF instrument using electrospray ionization (ESI, positive or negative ion mode).

**N-(1-(3-bromo-4-methylphenyl)ethyl)-4-chloro-3-(trifluoromethyl)aniline**

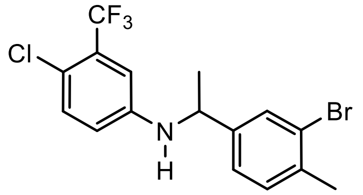


To a mixture of 1-(3-bromo-4-methylphenyl)ethan-1-one (1.11 g, 5.22 mmol), 4-chloro-3-(trifluoromethyl)aniline (1.02 g, 5.22 mmol) and methanol (10 mL) was added decaborane (191 mg, 1.56 mmol), and the mixture was stirred at room temperature overnight. After the mixture was diluted with ethyl acetate, aminopropyl silica gel powder was added. The resulting mixture was stirred at room temperature for 15 minutes and filtered. The filtrate was concentrated under reduced pressure. The residue was purified by column chromatography on aminopropyl silica gel (eluent : ethyl acetate/n-hexane = 0-15 %) to give the title compound (2.05 g, 100 %). $^{1}$H NMR ($CDCl_3$) δ: 1.51 (d, $J$ = 6.8 Hz, 3H), 2.37 (s, 3H), 4.15-4.27 (m, 1H), 4.33-4.44 (m, 1H), 6.48 (dd, $J$ = 8.8, 2.8 Hz, 1H), 6.83 (d, $J$ = 2.8 Hz, 1H), 7.10-7.23 (m, 3H), 7.49 (d, $J$ = 1.5 Hz, 1H).

**N-(1-(3-bromo-4-methylphenyl)ethyl)-4-chloro-3-methylaniline**

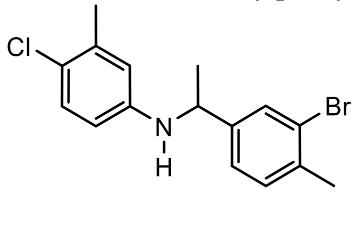

This title compound was prepared from 4-chloro-3-methylaniline, as described for the synthesis of N-(1-(3-bromo-4-methylphenyl)ethyl)-4-chloro-3-(trifluoromethyl)aniline, in 83 % yield. $^{1}$H NMR ($CDCl_3$) δ: 1.47 (d, $J$ = 6.8 Hz, 3H), 2.22 (s, 3H), 2.36 (s, 3H), 3.85-4.00 (m, 1H), 4.30-4.42 (m, 1H), 6.23 (dd, $J$ = 8.5, 2.8 Hz, 1H), 6.38 (d, $J$ = 2.8 Hz, 1H), 7.01 (d, $J$ = 8.5 Hz, 1H), 7.12-7.21 (m, 2H), 7.47-7.54 (m, 1H).

**N-(1-(3-bromo-4-methylphenyl)ethyl)-4-chloroaniline**

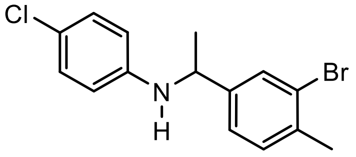


This title compound was prepared from 4-chloroaniline, as described for the synthesis of N-(1-(3-bromo-4-methylphenyl)ethyl)-4-chloro-3-(trifluoromethyl)aniline, in 92 % yield. $^{1}$H NMR ($CDCl_3$) δ: 1.48 (d, $J$ = 6.8 Hz, 3H), 2.36 (s, 3H), 3.95-4.08 (m, 1H), 4.30-4.43 (m, 1H), 6.35-6.44 (m, 2H), 6.94-7.07 (m, 2H), 7.11-7.21 (m, 2H), 7.46-7.53 (m, 1H).

**N-(1-(3-bromo-4-methylphenyl)ethyl)aniline**

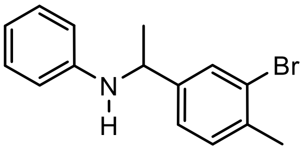


This title compound was prepared from aniline, as described for the synthesis of N-(1-(3-bromo-4-methylphenyl)ethyl)-4-chloro-3-(trifluoromethyl)aniline, in 100 % yield. $^{1}$H NMR ($CDCl_3$) δ: 1.49 (d, $J$ = 6.6 Hz, 3H), 2.36 (s, 3H), 3.92-4.05 (m, 1H), 4.41 (q, $J$ = 6.6 Hz, 1H), 6.45-6.53 (m, 2H), 6.62-6.70 (m, 1H), 7.05-7.13 (m, 2H), 7.14-7.23 (m, 2H), 7.51-7.55 (m, 1H).

**N-(1-(3-bromo-4-methylphenyl)ethyl)-4-chloro-N-ethyl-3-(trifluoromethyl)aniline**

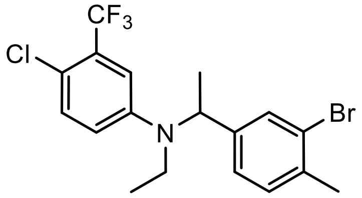


To a mixture of N-(1-(3-bromo-4-methylphenyl)ethyl)-4-chloro-3-(trifluoromethyl)aniline (1.11g, 2.83 mmol), acetaldehyde (5mol/L tetrahydrofuran solution, 1.7 mL, 8.49 mmol), tetrahydrofuran (3 mL) and methanol (3 mL) was added decaborane (173 mg, 1.41 mmol), and the mixture was stirred at room temperature for 1 hour. After the mixture was diluted with ethyl acetate, aminopropyl silica gel powder was added. The resulting mixture was stirred at room temperature for 15 minutes and filtered. The filtrate was concentrated under reduced pressure. The residue was purified by column chromatography on aminopropyl silica gel (eluent : n-hexane) to give the title compound (1.19 g, 100 %). $^1$H NMR ($CDCl_3$) δ: 1.08 (t, *J* = 7.0 Hz, 3H), 1.56 (d, *J* = 7.0 Hz, 3H), 2.38 (s, 3H), 3.24 (q, *J* = 7.0 Hz, 2H), 4.95 (q, *J* = 7.0 Hz, 1H), 6.78 (dd, *J* = 9.0, 3.0 Hz, 1H), 7.03 (d, *J* = 3.0 Hz, 1H), 7.06-7.10 (m, 1H), 7.18 (d, *J* = 7.7 Hz, 1H), 7.23-7.28 (m, 1H), 7.41-7.46 (m, 1H).

**N-(1-(3-bromo-4-methylphenyl)ethyl)-4-chloro-N-ethylaniline**

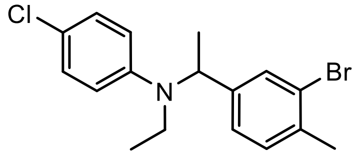


This title compound was prepared from N-(1-(3-bromo-4-methylphenyl)ethyl)-4-chloroaniline, as described for the synthesis of N-(1-(3-bromo-4-methylphenyl)ethyl)-4-chloro-N-ethyl-3-(trifluoromethyl)aniline, in 92 % yield. $^1$H NMR ($CDCl_3$) δ: 1.06 (t, *J* = 7.0 Hz, 3H), 1.53 (d, *J* = 6.9 Hz, 3H), 2.37 (s, 3H), 3.11-3.26 (m, 2H), 4.90 (q, *J* = 6.9 Hz, 1H), 6.63-6.75 (m, 2H), 7.05-7.21 (m, 4H), 7.41-7.48 (m, 1H).

**N-(1-(3-bromo-4-methylphenyl)ethyl)-N-ethylaniline**

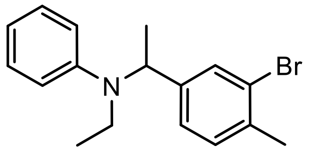


This title compound was prepared from N-(1-(3-bromo-4-methylphenyl)ethyl)aniline, as described for the synthesis of N-(1-(3-bromo-4-methylphenyl)ethyl)-4-chloro-N-ethyl-3-(trifluoromethyl)aniline, in 68 % yield. $^1$H NMR ($CDCl_3$) δ: 1.07 (t, *J* = 7.0 Hz, 3H), 1.53 (d, *J* = 6.9 Hz, 3H), 2.37 (s, 3H), 3.20 (q, *J* = 7.0 Hz, 2H), 4.98 (q, *J* = 6.9 Hz, 1H), 6.68-6.75 (m, 1H), 6.76-6.83 (m, 2H), 7.10-7.18 (m, 2H), 7.19-7.25 (m, 2H), 7.45-7.51 (m, 1H).

**N-(1-(3-bromo-4-methylphenyl)ethyl)-4-chloro-N-methyl-3-(trifluoromethyl)aniline**

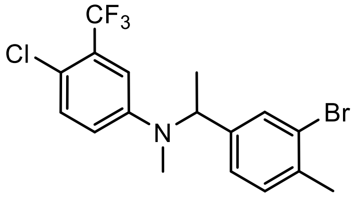


To a mixture of N-(1-(3-bromo-4-methylphenyl)ethyl)-4-chloro-3-(trifluoromethyl)aniline (0.306g, 0.779 mmol), formaldehyde (36 % aqueous solution, 0.20 mL, 2.34 mmol), tetrahydrofuran (1 mL) and methanol (2 mL) was added decaborane (48 mg, 0.390 mmol), and the mixture was stirred at room temperature overnight. After the mixture was diluted with ethyl acetate, aminopropyl silica gel powder was added. The resulting mixture was stirred at room temperature for 15 minutes and filtered. The filtrate was concentrated under reduced pressure. The residue was purified by column chromatography on aminopropyl silica gel (eluent : n-hexane) to give the title compound (0.268 g, 85 %). $^1$H NMR ($CDCl_3$) δ: 1.50-1.56 (m, 3H), 2.38 (s, 3H), 2.70 (s, 3H), 5.01 (q, *J* = 7.0 Hz, 1H), 6.83 (dd, *J* = 9.0, 3.0 Hz, 1H), 7.05 (d, *J* = 3.0 Hz, 1H), 7.06-7.10 (m, 1H), 7.19 (d, *J* = 7.9 Hz, 1H), 7.29 (d, *J* = 9.0 Hz, 1H), 7.42-7.45 (m, 1H).

**N-(1-(3-bromo-4-methylphenyl)ethyl)-4-chloro-N,3-dimethylaniline**

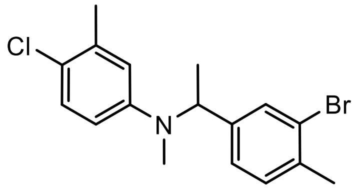


This title compound was prepared from N-(1-(3-bromo-4-methylphenyl)ethyl)-4-chloro-3-methylaniline, as described for the synthesis of N-(1-(3-bromo-4-methylphenyl)ethyl)-4-chloro-N-methyl-3-(trifluoromethyl)aniline, in 41 % yield. $^1$H NMR ($CDCl_3$) δ: 1.49 (d, *J* = 6.8 Hz, 3H), 2.33 (s, 3H), 2.38 (s, 3H), 2.63 (s, 3H), 4.99 (q, *J* = 6.8 Hz, 1H), 6.59 (dd, *J* = 8.8, 3.0 Hz, 1H), 6.67 (d, *J* = 3.0 Hz, 1H), 7.11 (dd, *J* = 7.8, 1.3 Hz, 1H), 7.13-7.20 (m, 2H), 7.46 (d, *J* = 1.3 Hz, 1H).

**1-(5-(1-((4-chloro-3-(trifluoromethyl)phenyl)(ethyl)amino)ethyl)-2-methylbenzyl)azetidine-3-carboxylic acid (*(RS)*-KSI-6666)**

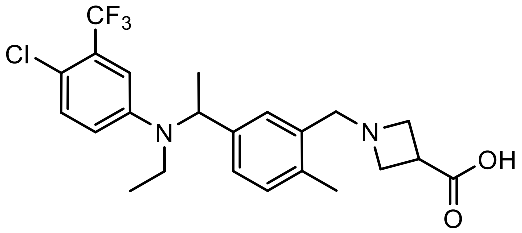


To a mixture of N-(1-(3-bromo-4-methylphenyl)ethyl)-4-chloro-N-ethyl-3-(trifluoromethyl)aniline (1.19 g, 2.83 mmol) and tetrahydrofuran (15 mL) was added n-butyllithium (2.65 mol/L hexane solution, 1.2 mL, 3.13 mmol) at -78 °C, and the mixture was stirred at same temperature for 30 minutes. To the reaction mixture was added N,N-dimethylformamide (1.1 mL, 14.1 mmol) at -78 °C, and the mixture was stirred at same temperature for 10 minutes. After the mixture was stirred at room temperature for 20 minutes, a saturated ammonium chloride aqueous solution was added. The resulting mixture was extracted with ethyl acetate. The extract was washed with brine and dried over anhydrous $MgSO_4$, and concentrated under reduced pressure. The residue was purified by column chromatography on silica gel (eluent : ethyl acetate/n-hexane = 0-10 %) to give 5-(1-((4-chloro-3-(trifluoromethyl)phenyl)(ethyl)amino)ethyl)-2-methylbenzaldehyde (652 mg, 62 %). This material (652 mg, 1.76 mmol) and azetidine-3-carboxylic acid methyl ester hydrochloride (669 mg, 4.41 mmol) were susupended in tetrahydrofuran (17 mL), and triethylamine (0.61 mL, 4.40 mmol) was added. After this mixture was stirred at room temperature for 15 minutes, sodium triacetoxyborohydride (1.87 g, 8.81 mmol) was added, and the mixture was stirred at room temperature overnight. After the mixture was diluted with ethyl acetate, the resulting mixture was washed with a saturated aqueous sodium hydrogen carbonate solution and brine successively. The organic layer was dried over anhydrous $MgSO_4$ and concentrated under reduced pressure. The residue was purified by column chromatography on aminopropyl silica gel (eluent : ethyl acetate/n-hexane = 0-35 %) to give methyl 1-(5-(1-((4-chloro-3-(trifluoromethyl)phenyl)(ethyl)amino)ethyl)-2-methylbenzyl)azetidine-3-carboxylate (628 mg, 76 %). This material (628 mg, 1.34 mmol) was dissolved in methanol (13 mL), and 2 mol/L sodium hydroxide solution (1.35 mL, 2.69 mmol) was added. The reaction mixture was stirred at room temperature overnight. 2 mol/L hydrochloric acid (1.35 mL, 2.69 mmol) was added to the reaction mixture, and concentrated under reduced pressure. The residue was diluted with water, and extracted by solid-phase extraction on a C18 Bond Elute cartride (eluent : methanol). The extract was concentrated under reduced pressure to give the title compound (574 mg, 94 %). $^{1}$H NMR ($CDCl_3$) δ: 1.03 (t, *J* = 7.0 Hz, 3H), 1.55 (d, *J* = 6.9 Hz, 3H), 2.37 (s, 3H), 3.10-3.32 (m, 3H), 3.76-3.89 (m, 2H), 4.00-4.25 (m, 4H), 4.97 (q, *J* = 6.9 Hz, 1H), 6.77 (dd, *J* = 9.0, 3.0 Hz, 1H), 7.00 (d, *J* = 3.0 Hz, 1H), 7.08-7.18 (m, 2H), 7.23 (d, *J* = 9.0 Hz, 1H), 7.28-7.35 (m, 1H); $^{13}$C NMR ($CDCl_3$) δ: 13.73, 17.45, 19.06, 34.69, 40.41, 55.98, 56.54, 56.63 (d, *J* = 1.5 Hz) 111.72 (q, *J* = 5.8 Hz), 117.00, 117.84 (d, *J* = 1.5 Hz), 123.19 (q, *J* = 273.2 Hz), 127.49, 128.34, 128.35 (q, *J* = 30.5 Hz), 130.01, 131.23, 131.88, 136.07, 140.40, 146.73, 176.10; HRMS-ESI (*m/z*): $[M + H]^+$ calcd for $C_{23}H_{27}ClF_3N_2O_2$, 455.1708; found 455.1720.

**1-(5-(1-((4-chlorophenyl)(ethyl)amino)ethyl)-2-methylbenzyl)azetidine-3-carboxylic acid (Compound 4)**

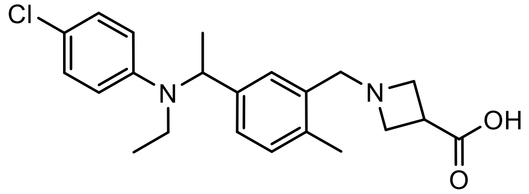


This title compound was prepared from N-(1-(3-bromo-4-methylphenyl)ethyl)-4-chloro-N-ethylaniline, as described for the synthesis of 1-(5-(1-((4-chloro-3-(trifluoromethyl)phenyl)(ethyl)amino)ethyl)-2-methylbenzyl)azetidine-3-carboxylic acid, in 70 % yield. $^{1}$H NMR ($CDCl_3$) δ: 1.02 (t, *J* = 7.0 Hz, 3H), 1.53 (d, *J* = 6.9 Hz, 3H), 2.37 (s, 3H), 3.08-3.28 (m, 3H), 3.77-3.94 (m, 2H), 4.00-4.25 (m, 4H), 4.91 (q, *J* = 6.9 Hz, 1H), 6.58-6.74 (m, 2H), 7.00-7.20 (m, 4H), 7.25-7.36 (m, 1H); $^{13}$C NMR ($CDCl_3$) δ: 13.99, 17.59, 19.10, 34.61, 40.47, 55.80, 56.41, 56.78, 115.19, 121.33, 127.83, 128.63, 128.91, 129.31, 131.18, 135.87, 141.28, 146.96, 175.76; HRMS-ESI (*m/z*): $[M + H]^+$ calcd for $C_{22}H_{28}ClN_2O_2$, 387.1834; found 387.1847.

**1-(5-(1-(ethyl(phenyl)amino)ethyl)-2-methylbenzyl)azetidine-3-carboxylic acid (Compound 5)**

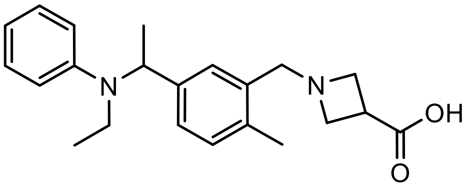


This title compound was prepared from N-(1-(3-bromo-4-methylphenyl)ethyl)-N-ethylaniline, as described for the synthesis of 1-(5-(1-((4-chloro-3-(trifluoromethyl)phenyl)(ethyl)amino)ethyl)-2-methylbenzyl)azetidine-3-carboxylic acid, in 48 % yield. $^{1}$H NMR ($CDCl_3$) δ: 1.04 (t, *J* = 7.0 Hz, 3H), 1.54 (d, *J* = 7.0 Hz, 3H), 2.36 (s, 3H), 3.06-3.30 (m, 3H), 3.67-3.87 (m, 2H), 3.92-4.24 (m, 4H), 4.99 (q, *J* = 7.0 Hz, 1H), 6.60-6.84 (m, 3H), 7.00-7.35 (m, 5H); $^{13}$C NMR ($CDCl_3$) δ: 14.21, 17.58, 19.04, 34.89, 40.28, 56.04, 56.45, 56.58, 114.02, 116.72, 127.87, 128.83, 129.17, 129.66, 131.06, 135.78, 141.51, 148.41, 176.13; HRMS-ESI (*m/z*): $[M + H]^+$ calcd for $C_{22}H_{29}N_2O_2$, 353.2224; found 353.2243.

**1-(5-(1-((4-chloro-3-(trifluoromethyl)phenyl)(methyl)amino)ethyl)-2-methylbenzyl)azetidine-3-carboxylic acid (Compound 3)**

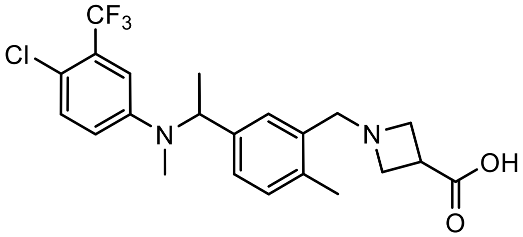


This title compound was prepared from N-(1-(3-bromo-4-methylphenyl)ethyl)-4-chloro-N-methyl-3-(trifluoromethyl)aniline, as described for the synthesis of 1-(5-(1-((4-chloro-3-(trifluoromethyl)phenyl)(ethyl)amino)ethyl)-2-methylbenzyl)azetidine-3-carboxylic acid, in 46 % yield. $^{1}$H NMR ($CDCl_3$) δ: 1.52 (d, *J* = 6.8 Hz, 3H), 2.37 (s, 3H), 2.66 (s, 3H), 3.08-3.24 (m, 1H), 3.72-3.89 (m, 2H), 4.00-4.25 (m, 4H), 5.02 (q, *J* = 6.8 Hz, 1H), 6.82 (dd, *J* = 9.0, 3.0 Hz, 1H), 7.03 (d, *J* = 3.0 Hz, 1H), 7.09-7.20 (m, 2H), 7.23-7.35 (m, 2H); $^{13}$C NMR ($CDCl_3$) δ: 16.40, 19.07, 32.08, 34.66, 56.04, 56.36, 56.70 (d, *J* = 2.2 Hz), 111.25 (q, *J* = 5.8 Hz), 116.51, 118.16 (d, *J* = 2.2 Hz), 123.18 (q, *J* = 273.2 Hz), 127.33, 128.16, 128.37 (q, *J* = 30.5 Hz), 130.02, 131.28, 131.90, 136.09, 140.15, 148.38, 175.95; HRMS-ESI (*m/z*): $[M + H]^+$ calcd for $C_{22}H_{25}ClF_3N_2O_2$, 441.1551; found 441.1565.

**1-(5-(1-((4-chloro-3-methylphenyl)(methyl)amino)ethyl)-2-methylbenzyl)azetidine-3-carboxylic acid (Compound 2)**

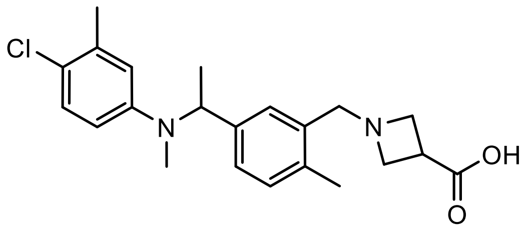

This title compound was prepared from N-(1-(3-bromo-4-methylphenyl)ethyl)-4-chloro-N,3-dimethylaniline, as described for the synthesis of 1-(5-(1-((4-chloro-3-(trifluoromethyl)phenyl)(ethyl)amino)ethyl)-2-methylbenzyl)azetidine-3-carboxylic acid, in 41 % yield. $^{1}$H NMR ($CDCl_3$) δ: 1.47 (d, *J* = 6.9 Hz, 3H), 2.30 (s, 3H), 2.36 (s, 3H), 2.57 (s, 3H), 3.12-3.30 (m, 1H), 3.75–3.96 (m, 2H), 4.00-4.22 (m, 4H), 5.00 (q, *J* = 6.8 Hz, 1H), 6.57 (dd, *J* = 8.8, 3.0 Hz, 1H), 6.65 (d, *J* = 3.0 Hz, 1H), 7.08-7.20 (m, 3H), 7.27-7.32 (m, 1H); $^{13}$C NMR ($CDCl_3$) δ: 16.04, 19.07, 20.59, 31.95, 34.78, 56.06, 56.43, 56.76 (d, *J* = 5.8 Hz), 112.16, 115.57, 121.97, 127.61, 128.54, 129.32, 129.72, 131.15, 135.92, 136.27, 140.82, 148.87, 176.10; HRMS-ESI (*m/z*): $[M + H]^+$ calcd for $C_{22}H_{28}ClN_2O_2$, 387.1834; found 387.1851.

**1-(3-(((4-chloro-3-methylphenyl)(methyl)amino)methyl)benzyl)azetidine-3-carboxylic acid (Compound 1)**

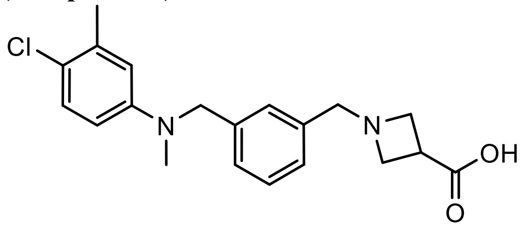

To a mixture of azetidine-3-carboxylic acid methyl ester hydrochloride (400 mg, 2.64 mmol), benzene-1,3-dicarbaldehyde (708 mg, 5.28 mmol), triethylamine (0.37 mL, 2.64 mmol) and tetrahydrofuran (10 mL) was added sodium triacetoxyborohydride (1.12 g, 5.28 mmol), and the mixture was stirred at room temperature for 3 hours. After the mixture was diluted with ethyl acetate, the resulting mixture was washed with a saturated aqueous sodium hydrogen carbonate solution and brine successively. The organic layer was dried over anhydrous $Na_2SO_4$ and concentrated under reduced pressure. The residue was purified by column chromatography on silica gel (eluent : ethyl acetate/n-hexane = 20-100 %) to give methyl 1-(3-formylbenzyl)azetidine-3-carboxylate (330 mg, 54 %). This material (330 mg, 1.42mmol) and 4-chloro-3-methylaniline (243 mg, 1.72mmol) were dissolved in methanol (8 mL), and decaborane (86 mg, 0.704 mmol) was added. The mixture was stirred at room temperature for 2 hours. After the mixture was diluted with ethyl acetate, aminopropyl silica gel powder was added. The resulting mixture was stirred at room temperature for 15 minutes and filtered. The filtrate was concentrated under reduced pressure. The residue was purified by column chromatography on aminopropyl silica gel (eluent : ethyl acetate/n-hexane = 20-50 %) to give methyl 1-(3-(((4-chloro-3-methylphenyl)amino)methyl)benzyl)azetidine-3-carboxylate

(451 mg, 89 %). This material (266 mg, 0.741 mmol) and formaldehyde solution (37 % aqueous solution, 0.35 mL, 4.31 mmol) were dissolved in methanol (5 mL), and decaborane (45 mg, 0.368 mmol) was added. The mixture was stirred at room temperature for 1 hour. After the mixture was diluted with ethyl acetate, aminopropyl silica gel powder was added. The resulting mixture was stirred at room temperature for 15 minutes and filtered. The filtrate was concentrated under reduced pressure. The residue was purified by column chromatography on silica gel (eluent : ethyl acetate/n-hexane = 50-100 %) to give methyl 1-(3-(((4-chloro-3-methylphenyl)(methyl)amino)methyl)benzyl)azetidine-3-carboxylate (162 mg, 59 %). This material (137 mg, 0.367 mmol) was dissolved in 1,4-dioxane (2 mL) and methanol (2 mL), and 2 mol/L sodium hydroxide solution (0.55 mL, 1.10 mmol) was added. The mixture was stirred at room temperature for 2 hours. 2 mol/L hydrochloric acid (0.55 mL, 1.10 mmol) was added, and the resulting mixture was concentrated under reduced pressure. The residue was diluted with ethyl acetate, and washed with water. The organic layer was concentrated under reduced pressure to give the title compound (88 mg, 67 %). $^1$H NMR ($CDCl_3$) δ: 2.28 (s, 3H), 2.96 (s, 3H), 3.22-3.40 (m, 1H), 3.80-4.25 (m, 6H), 4.47 (s, 2H), 6.46 (dd, $J$ = 8.8, 3.0 Hz, 1H), 6.56 (d, $J$ = 3.0 Hz, 1H), 7.02-7.39 (m, 5H) ; $^{13}$C NMR ($CDCl_3$) δ: 20.56, 34.35, 38.77, 56.04, 56.49, 58.57, 111.40, 114.73, 122.07, 127.73, 128.15, 128.41, 129.36, 129.52, 130.75, 136.33, 140.01, 148.29, 175.66; HRMS-ESI (*m/z*): $[M + H]^+$ calcd for $C_{20}H_{24}ClN_2O_2$, 359.1521; found 359.1524.

***(R)*-1-(5-(1-((4-chloro-3-(trifluoromethyl)phenyl)(ethyl)amino)ethyl)-2-methylbenzyl)azetidine-3-carboxylic acid (*(R)*-KSI-6666)**

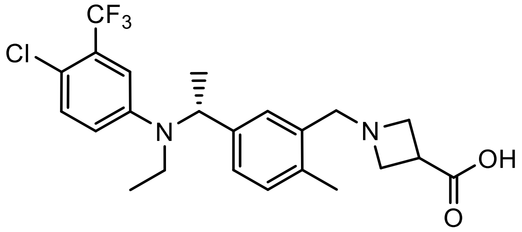


methyl 1-(5-(1-((4-chloro-3-(trifluoromethyl)phenyl)(ethyl)amino)ethyl)-2-methylbenzyl)azetidine-3-carboxylate (1.69 g, 3.60mmol) was separated by an optically active column chromatography (column : CHIRALCEL OJ-3 (4.6 mm x 250 mm), eluent : n-hexane/2-propanol/diethylamine = 95/5/0.1, flow rate : 1.0 mL/min, temp. : 40 ℃) to give methyl *(R)*-1-(5-(1-((4-chloro-3-(trifluoromethyl)phenyl)(ethyl)amino)ethyl)-2-methylbenzyl)azetidine-3-carboxylate* (700 mg, 42 %, retention time 7.2 min). This material (500 mg, 1.07 mmol) was dissolved in methanol (10 mL), and 2 mol/L sodium hydroxide solution (1.07 mL, 2.14 mmol) was added. The reaction mixture was stirred at room temperature overnight. 2 mol/L hydrochloric acid (1.07 mL, 2.14 mmol) was added to the reaction mixture, and concentrated under reduced pressure. The residue was diluted with water, and extracted by solid-phase extraction on a C18 Bond Elute cartride (eluent : methanol). The extract was concentrated under reduced pressure to give the title compound (449 mg, 93 %).* The absolute configuration was determined by comparison with methyl *(R)*-1-(5-(1-((4-chloro-3-(trifluoromethyl)phenyl)(ethyl)amino)ethyl)-2-methylbenzyl)azetidine-3-carboxylate derived from a chiral amine. $^1$H NMR ($CDCl_3$) δ: 1.03 (t, *J* = 7.0 Hz, 3H), 1.55 (d, *J* = 6.9 Hz, 3H), 2.37 (s, 3H), 3.10-3.32 (m, 3H), 3.76-3.89 (m, 2H), 4.00-4.25 (m, 4H), 4.97 (q, *J* = 6.9 Hz, 1H), 6.76 (dd, *J* = 9.0, 3.0 Hz, 1H), 7.00 (d, *J* = 3.0 Hz, 1H), 7.08-7.18 (m, 2H), 7.23 (d, *J* = 9.0 Hz, 1H), 7.28-7.35 (m, 1H); $^{13}$C NMR ($CDCl_3$) δ: 13.73, 17.47, 19.06, 34.72, 40.41, 56.12, 56.56, 56.65 (d, *J* = 1.5 Hz), 111.72 (q, *J* = 5.8 Hz), 116.99, 117.84 (d, *J* = 1.5 Hz), 123.19 (q, *J* = 273.2 Hz), 127.42, 128.29, 128.36 (q, *J* = 30.5 Hz), 130.21, 131.20, 131.87, 136.04, 140.36, 146.74, 176.08; HRMS-ESI (*m/z*): $[M + H]^+$ calcd for $C_{23}H_{27}ClF_3N_2O_2$, 455.1708; found 455.1728.

***(S)*-1-(5-(1-((4-chloro-3-(trifluoromethyl)phenyl)(ethyl)amino)ethyl)-2-methylbenzyl)azetidine-3-carboxylic acid (*(S)*-KSI-6666)**

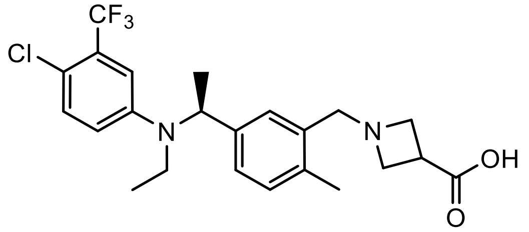


methyl 1-(5-(1-((4-chloro-3-(trifluoromethyl)phenyl)(ethyl)amino)ethyl)-2-methylbenzyl)azetidine-3-carboxylate (1.69 g, 3.60mmol) was separated by an optically active column chromatography (column : CHIRALCEL OJ-3 (4.6 mm x 250 mm), eluent : n-hexane/2-propanol/diethylamine = 95/5/0.1, flow rate : 1.0 mL/min, temp. : 40 °C) to give methyl *(S)*-1-(5-(1-((4-chloro-3-(trifluoromethyl)phenyl)(ethyl)amino)ethyl)-2-methylbenzyl)azetidine-3-carboxylate (700 mg, 42 %, retention time 6.1 min). This material (496 mg, 1.06 mmol) was dissolved in methanol (10 mL), and 2 mol/L sodium hydroxide solution (1.07 mL, 2.14 mmol) was added. The reaction mixture was stirred at room temperature for 2 hours. 2 mol/L hydrochloric acid (1.07 mL, 2.14 mmol) was added to the reaction mixture, and concentrated under reduced pressure. The residue was suspended with water (10 mL) and extracted with 10% methanol/ ethyl acetate solution in twice. The organic layer was washed with brine and dried over anhydrous $MgSO_4$, and concentrated under reduced pressure to give the title compound (465 mg, 97 %). $^1$H NMR ($CDCl_3$) δ: 1.03 (t, *J* = 7.0 Hz, 3H), 1.56 (d, *J* = 6.8 Hz, 3H), 2.36 (s, 3H), 3.12-3.50 (m, 3H), 3.76-4.40 (m, 6H), 4.96 (q, *J* = 6.8 Hz, 1H), 6.77 (dd, *J* = 9.0, 2.9 Hz, 1H), 6.98 (d, *J* = 2.9 Hz, 1H), 7.08-7.18 (m, 2H), 7.22 (d, *J* = 9.0 Hz, 1H), 7.30-7.47 (m, 1H); $^{13}$C NMR ($CDCl_3$) δ: 13.73, 17.59, 19.16, 34.72, 40.51, 55.69, 56.53, 56.59, 111.74 (q, *J* = 5.8 Hz), 117.08, 117.85, 123.18 (q, *J* = 273.2 Hz), 127.80, 128.30 (q, *J* = 30.5 Hz), 128.61, 129.27, 131.31, 131.88, 136.23, 140.62, 146.69, 176.34; HRMS-ESI (*m/z*): $[M + H]^+$ calcd for $C_{23}H_{27}ClF_3N_2O_2$, 455.1708; found 455.1723.
